# Plant sexual and asexual contributions to the seed microbiome

**DOI:** 10.1101/2025.06.16.659819

**Authors:** Maria Faticov, Ayco J. M. Tack, Doris Ortner, Gabriele Berg, Ahmed Abdelfattah

**Affiliations:** Department of Wildlife, Fish and Environmental Studies, SLU, Umeå, Sweden; Department of Ecology, Environment and Plant Sciences, SU, Stockholm, Sweden; Leibniz Institute for Agricultural Engineering and Bioeconomy (ATB), Potsdam, Germany; Institute for Biochemistry and Biology, University of Potsdam, Potsdam, Brandenburg, Germany; Institute of Environmental Biotechnology, Graz University of Technology, Graz, Styria, Austria

**Keywords:** bacteria, community assembly, flower, microbial transmission, seed microbiome

## Abstract

The seed microbiome plays a key role in the assembly of the plant microbiome, which has major impacts on plant functioning. Nonetheless, little is known about the origin of the seed microbiome. We investigated the relative contributions of two hypothesized transmission routes: sexual inheritance (via reproductive organs) and asexual inheritance (via the plant vascular system). To do that, we sampled shoot endophytes both before bloom and at seed maturity stages, apple flower ovaries and pollen sacs, and mature seeds from field-grown apple trees (*Malus domestica* Borkh. cv ‘Gala Galaxy Selecta’). We showed that bacterial richness, diversity and composition differ among tissue types, with shoots before bloom harboring a higher diversity than ovary and pollen. Source tracking revealed that both sexual (30.3%) and asexual (23.8%) pathways contributed to seed microbiome assembly, with shoots at seed maturity being the dominant source. Notably, a large proportion (49.5%) of the seed microbiome originated from unknown sources. Lastly, the transmission pathways significantly differed among bacterial genera, with *Pseudomonas* primarily linked to shoots, *Rhizobacter* to pollen and *Burkholderia* to the ovary. Insights into seed microbiome origin offers new opportunities to enhance seed health and crop productivity through microbiome-assisted breeding.

## Introduction

The concept of phylosymbiosis postulates that the similarity between host-associated microbial communities is mirrored by the evolutionary relationships among host (Brucker & Bordenstein, 2013). It suggests that closely related plant species harbor more similar microbiomes than distantly related species due to shared evolutionary history (Brooks *et al*., 2016; Lim & Bordenstein, 2020). Although phylosymbiosis can arise through various mechanisms, including coevolution, shared environmental factors, or host selection effects (Kohl, 2020) one mechanism that is commonly overlooked is microbial inheritance. Microbial inheritance is the process in which plant-associated microorganisms pass from parents to seed, and persist seed dormancy and germination (Abdelfattah *et al*., 2023). By transmission through the seeds, inherited microorganisms can be maintained across generations, preserving part of the plant microbiome over time (Abdelfattah *et al*., 2023; Sulesky-Grieb *et al*., 2024). For this mechanism to contribute to phylosymbiosis, however, it would require that introgression events not only include genetic material but also microbial communities from each parent plant, which in turn would lead to a novel admixture of both partners. Yet, seed microbial communities are not solely shaped by inherited microorganisms. For example, part of the seed microbes may originate from environmental sources such as air, water, and pollinators (Prado *et al*., 2020; Simonin *et al*., 2022). Understanding the role of microbial inheritance is essential for evaluating the evolutionary role of seed microbiomes - and by extension, for testing whether microbial inheritance can reinforce phylosymbiotic patterns.

The transmission of microbes from plant to seed can occur via two main pathways: sexual, involving reproductive organs such as pollen and ovaries (Mitter *et al*., 2017; Bergmann *et al*., 2025) and asexual, originating from vegetative tissues like shoots (Abdelfattah *et al*., 2021, 2023). For example, if the seed microbiome is predominantly originated from the sexual transmission pathway, one can expect that the microbial communities in shoots before bloom would also be important sources of microbes (Olimi *et al*., 2022). This is because floral organs develop directly from these shoots, and at the time of flowering, shoot endophytes are likely the primary microbial source for emerging reproductive tissues (Abdelfattah *et al*., 2023; War *et al*., 2023). In other words, any microbes transmitted through pollen or ovary most likely originate from or pass through vegetative tissues. Conversely, if the seed microbiome is primarily shaped by asexual transmission, then shoots collected during seed maturation would be more important sources of the seed microbiome, as they remain connected through vascular system to the maturing seed and can enable microbial transfer through maternal tissues during seed filling. These two scenarios are not mutually exclusive, but each pathway implies distinct temporal and spatial patterns of microbial transmission toward the seed. Understanding the relative contributions of sexual and asexual transmission pathways will not only clarify the mechanisms underlying microbial inheritance but also provide insights into the ecological and evolutionary forces shaping seed microbiomes. This knowledge can enhance our understanding of host-microbe coevolution and phylosymbiotic patterns within plant lineages.

Microbial communities are not homogeneously distributed within plants. Instead, different tissues such as stems, leaves, flowers, and reproductive organs tend to harbor distinct microbiomes shaped by distinct physiological characteristics (Hardoim et al., 2015; Compant et al., 2011). Studies have shown that pollen grains can carry diverse microorganisms that could be transmitted during fertilization (Khalaf *et al*., 2023; Cardinale & Schnell, 2024; Armstrong *et al*., 2024). Paternal breeding lines contributed substantially to the seed microbiota of oilseed rape (Wassermann *et al*., 2022). Although the presence of microorganisms in the plant ovules was reported (Mundt & Hinkle, 1976) to this date much less is known about female than male reproductive parts (Bergmann *et al*., 2025). As for asexual tissues, shoots and stems often support more diverse microbial communities than reproductive tissues, likely due to their extended developmental windows and greater exposure to microbes in the environment. Interestingly, transmission pathways may differ among microbial taxa, with certain genera associated with specific tissues or transmission route (Chesneau *et al*., 2020). Altogether it is important to consider both tissue-specific microbes and taxon-specific transmission strategies when studying plant-to-seed microbial inheritance.

This study aims to address the critical knowledge gap regarding the origins of the seed microbiome. Specifically, we aim to disentangle the contributions of sexual (pollen sacs and ovaries) and asexual (shoot endophytes) pathways to seed microbiome origin and assembly (Fig. 1). We hypothesize that:

**Fig. 1.**
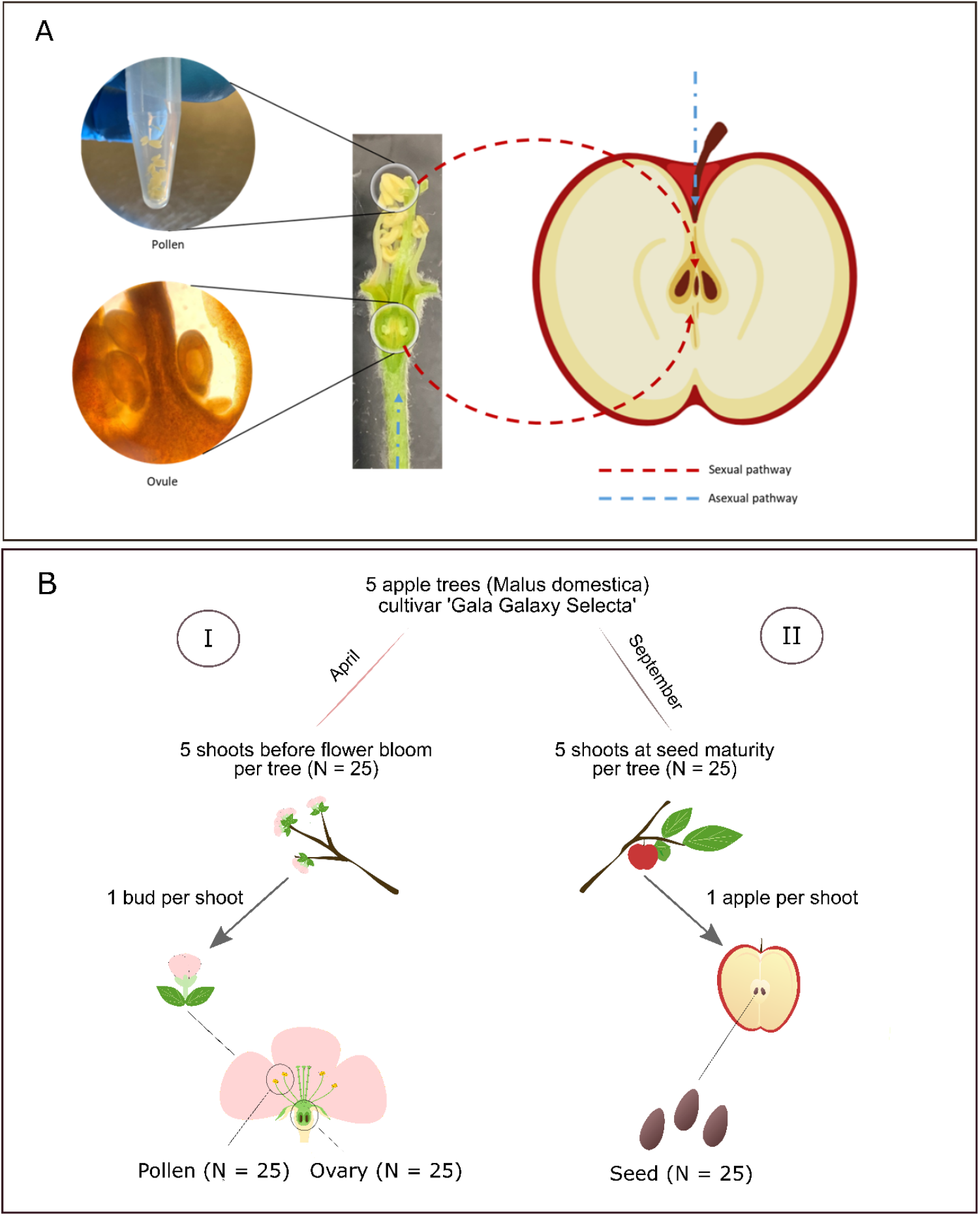
Schematic representation of microbial transmission pathways to the seed in apple trees. (A) Dashed orange arrows indicate the sexual transmission pathway, where microbes are transferred via floral organs including pollen and ovary tissues. Dashed blue arrows indicate the asexual transmission pathway, through which microbes colonize seeds via shoot tissues both before flower bloom and during seed maturity. (B) Experimental design showing the sampling scheme. From each of five ‘Gala Galaxy Selecta’ apple trees, we collected samples from five shoots (I) prior to flowering (April) and (II) at seed maturity (September). From the five shoots prior to flowering, we collected shoot tissue and five floral buds. In the laboratory, each flower bud was dissected to isolate ovary and pollen. From the five shoots at seed maturation, we collected shoot tissues, five apples and seeds. A total of 25 replicate samples were collected for each tissue type.

1. Microbial diversity and composition differ between shoots, ovary and pollen, with:
  a. Shoots harboring a higher diversity of microorganisms than the ovary and pollen.
  b. Shoot endophytes before bloom and at seed maturity are compositionally more similar to each other than to the ovary and pollen tissues.
2. Both sexual (ovary and pollen) and asexual (shoots before bloom and at seed maturity) pathways are hypothesized to contribute to seed microbiome assembly, though their relative importance can vary. Thus, we expected that:
  a. If sexual transmission dominates, shoots at bloom can serve as the main source for microbes colonizing emerging floral organs, making shoot at bloom, ovaries, and pollen sacs similar in their community composition to seed
  b. If asexual transmission dominates, we expect shoots at seed maturity to contribute the largest proportion to the seed microbiome, due to microbial transmission through the vascular system to developing seeds
3. Some microbial taxa are predominantly associated with either asexual (e.g., *Allorhizobium* and *Pseudomonas* from shoots) or sexual (e.g., *Rhizobacter* from pollen and *Erwinia* from ovary or pollen) tissues.
4. A notable proportion of the seed microbiome originates from unknown sources, possibly reflecting unsampled environmental reservoirs such as pollinators or air.

## Materials and Methods

### Study system and sample collection

To study the vertical transmission of endophytes in plants, we selected five apple trees (*Malus domestica*, cultivar ‘Gala Galaxy Selecta’) from the Botanical Garden of the University of Graz (47.0814° N, 15.4562° E). Sampling was conducted at two time points: before bloom (April 15, 2020) and at seed maturity (September 9, 2020) (Fig. 1b). All samples were collected using sterile forceps and gloves, transported on ice and processed under sterile conditions in a laminar flow cabinet.

To investigate the floral community, five flower buds from each of five trees were collected right before tree bloom in April (Fig. 1a). Alongside flower buds, shoots leading to the sampled flowers were collected at the same time. In September, mature apples were harvested, and seeds were extracted under sterile lab conditions (Fig. 1a). Alongside seeds, shoots leading to the sampled fruits were collected at the same time. The final set of samples included 25 pollen, 25 ovary and 25 shoot samples during bloom and 25 seed and 25 shoot samples during seed maturity (Fig. 1b).

### Preparation of ovary, pollen, shoot and seed samples

Under sterile laboratory conditions, each flower bud was dissected to isolate the female (ovary) and male (pollen sac) reproductive organs (Fig. 1a). The outer ovary layer was removed, and the entire inner portion was processed as a single ovary sample. Closed pollen sacs (hereafter referred to as *pollen*) were isolated from the same buds, with all pollen sacs from a single bud pooled into a single sample. Samples were stored at −20°C until DNA extraction.

Shoots collected before bloom and during seed maturity were surface sterilized in a 4% Sodium Hypochlorite (NaOCl) solution then rinsed three times with autoclaved Ultraopure water, each rinse lasting three minutes. After sterilization, the bark of the shoots was removed using a scalpel, and the first and last centimeters of the shoots were discarded. The shoots were then cut into 1 cm pieces, weighed, and transferred into sterile bags. To prepare the samples, 10 mL of 0.85% sodium chloride (NaCl) solution was added, and the tissue was homogenized using a sterilized mortar and pestle. The homogenate was transferred into two Eppendorf tubes and centrifuged at 16,000 g for 20 minutes at 4°C. After discarding the supernatant, 2 mL of NaCl solution was added to each tube for a second centrifugation under the same conditions. Seeds were processed using the same protocol. For each seed sample, approximately 8 seeds were ground in 10 mL of 0.85% NaCl solution, followed by centrifugation steps. All resulting pellets (from seeds and shoots) were stored at −20°C until DNA extraction.

### Molecular methods and bioinformatics

DNA extraction was performed using the FastDNA™ SPIN Kit for Soil (MP Biomedicals). DNA extractions were carried out separately for each sample type (ovules, pollen, seeds, shoots before bloom and at seed maturity) to prevent microbial cross-contamination, particularly between pollen and ovule samples. The 16S rRNA gene was amplified using primers 515f and 806r (Walters *et al*., 2015). To minimize host mitochondrial and plastid 16S rDNA amplification, Peptide Nucleic Acid (PNA) clamps were added to the PCR mixture (von Wintzingerode *et al*., 2000; Lundberg *et al*., 2013). PCR reactions were performed in a total volume of 30 µL and were purified using the Wizard SV Gel and PCR Clean-Up System (Promega, Madison, WI, USA; see Supporting Information for details). DNA concentration was measured with a Nanodrop 2000 spectrophotometer (Thermo Fisher Scientific, Wilmington, DE, USA), samples were pooled at equimolar concentrations and sequenced using 250 bp paired-end Illumina MiSeq sequencer.

Raw sequence data were processed using QIIME2 (version 2021.2) (Caporaso *et al*., 2010; Bolyen *et al*., 2019). Following paired-end read joining and quality filtering with Phred-score 33, error correction, denoising, chimera removal, and Amplicon Sequence Variant (ASV) table generation were performed using DADA2 (Callahan *et al*., 2016). Taxonomic assignment of ASVs was conducted using the SILVA reference database (version 132, 99% identity, 06.05.21), with VSEARCH implemented for de novo clustering. Sequences of host mitochondrial and chloroplast origin were filtered out, and ASVs detected in negative controls were removed for each apple tissue type independently.

### Statistical analysis

All analyses were conducted using R (version 4.4.3) (R Core Team, 2022). To investigate bacterial community composition, species richness, evenness, and diversity (Shannon index), the ASV table was rarefied to 200 reads per sample. Pairwise comparisons of microbial richness, diversity and evenness among tissues, such as ovaries, pollen, shoots and seeds, were assessed using Tukey’s post hoc test. Cumulative Sum Scaling (CSS) from the *MetagenomeSeq* package was applied to account for differences in sequencing depth (Paulson *et al*., 2013). Beta diversity was calculated using Bray-Curtis dissimilarity metrics (Anderson, 2001; McArdle & Anderson, 2001), and multivariate microbial community composition was visualized through Principal Coordinates Analysis (PCoA) using *plot_ordination* function in the *ggplot2* package (Wickham, 2009). Statistical testing of beta diversity was performed using PERMANOVA with the *adonis2* function in the *vegan* package (Oksanen *et al*., 2022). Finally, to assess whether microbial communities in shoots before bloom were more similar to those at seed maturity than to those in ovary or pollen tissues, we calculated Bray–Curtis dissimilarities between sample pairs from shoots at bloom and each of the following: shoot at seed maturity, ovary, and pollen. Distances were extracted from the dissimilarity matrix and grouped by tissue pair. We then performed pairwise Wilcoxon rank-sum tests to compare the distributions of these between-group distances.

To infer the relative contributions of sexual (ovaries and pollen) and asexual (shoot) tissues to seed microbiome assembly, we applied two complementary source-tracking approaches. First, we used a Bayesian source-tracking method implemented in SourceTracker v2.0.1 (Knights *et al*., 2011), which identifies taxa that differ among sources and sinks. Second, we applied FEAST v0.1.0 (Shenhav et al., 2019) with a maximum of 5000 iterations. To enhance the robustness of our findings, we applied both methods, as they offer complementary strengths. SourceTracker provides taxon-specific insights by identifying differences among sources and sinks, while FEAST is computationally efficient but lacks taxonomic resolution. In both source tracking approaches, seeds were set as the “sink” samples, while ovary, pollen, shoot samples before bloom and at seed maturity were defined as “source”. The algorithm identified an unexplained fraction, named ‘unknown’ source, which represented ASVs found in the sink, but originating from other unsampled sources. Since both methods showed quantitatively and qualitatively similar results, we present SourceTracker results in the main text, while FEAST results are presented in the supplementary material (Fig. S1-S3).

## Results

The bacterial taxa with the highest relative abundance varied across tissues. In ovaries, the dominant genera were *Burkholderia, Enhydrobacter*, and *Ralstonia*, followed by *Pseudomonas* (Fig. 2). In pollen, the dominant genera were *Burkholderia, Pseudomonas, Candidatus*, and *Rhizobacter* (Fig. 2). For shoots, the composition shifted with developmental stage: before bloom, *Novosphingobium* and *Allorhizobium* were the most prevalent, whereas at seed maturity, *Burkholderia, Allorhizobium*, and *Enhydrobacter* dominated (Fig. 2). Finally, in seeds, *Burkholderia* and *Pseudomonas* were the most abundant bacterial genera (Fig. 2).

**Fig. 2.**
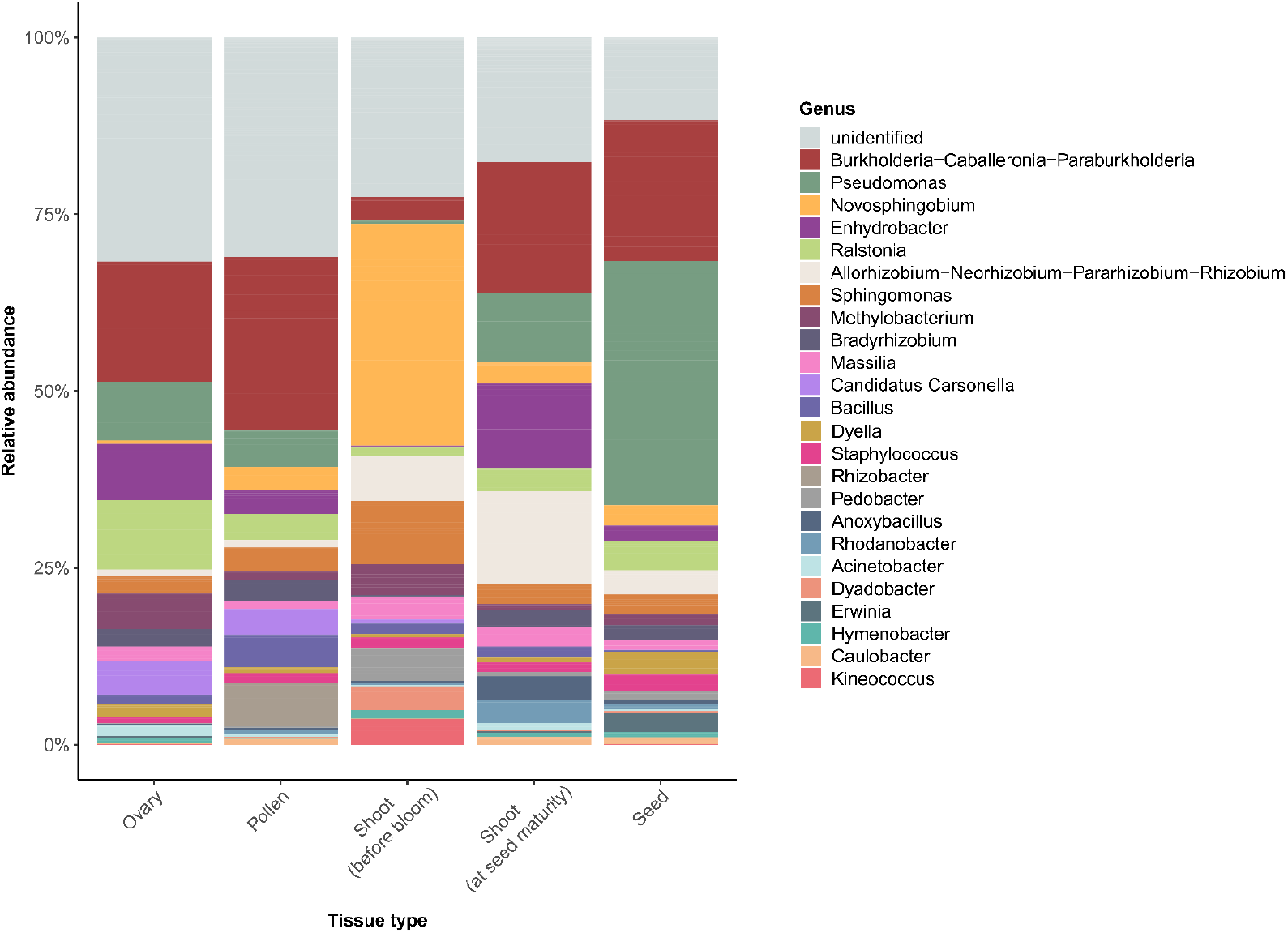
Relative abundance of bacterial genera in ovary, pollen, shoots before bloom and shoots at seed maturity. Shown are the 25 genera with the highest relative abundance.

*The richness, diversity and composition of the shoot, ovary, pollen and seed microbiomes* Bacterial richness varied significantly among tissue types, with the lowest richness in the ovary and seed, intermediate richness in pollen, and the highest richness in shoots (F4, 97 = 9.97, p < 0.0001). Post-hoc comparisons revealed that the ovary had significantly lower richness than pollen, shoots before bloom, and shoots at seed maturity, but was equal in richness to seeds (Fig. 3a; Table S1a). Moreover, bacterial richness in seeds was significantly lower than in shoots before bloom (Fig. 3a; Table S1a). A comparable pattern emerged for Shannon diversity, which also varied significantly across tissue types (F4, 97 = 9.63, p < 0.0001; Fig. 3b). As for richness, post-hoc tests showed that ovary samples had significantly lower diversity than pollen, shoots before bloom, and shoots at seed maturity, but was equal in richness to seeds (Fig. 3b; Table S1b). Moreover, pollen and shoots at bloom had significantly higher diversity than seeds (Fig. 3b; Table S1b).

**Fig. 3.**
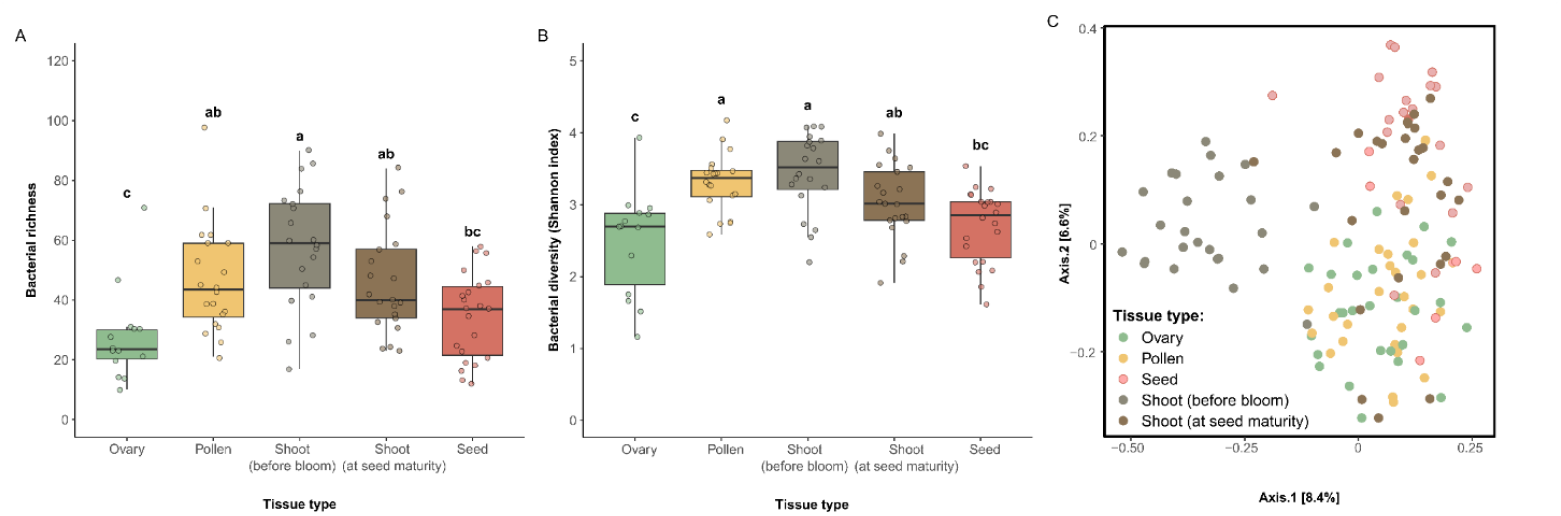
The differences in bacterial richness (a), diversity (b) and community composition among ovary, pollen, shoots before bloom, shoots at seed maturity and seed. (a-b) Differences in species richness and diversity among tissue types. The horizontal line within each box represents the median, the box edges show the interquartile range (IQR), and the whiskers demonstrate the data range. The small circles represent raw data points, which are horizontally jittered to avoid overlap. (c) Difference in community composition among tissue types visualized with two-dimensional principal coordinates analysis (PCoA) of bacterial communities based on Bray–Curtis dissimilarity index.

Bacterial community composition varied significantly across tissue types (R^2^ = 0.14, F = 5.30, p = 0.001; Fig. 3c), with all pairwise comparisons showing a significant differences (Table S2). The most pronounced compositional difference was observed between shoot endophytes before bloom and seeds (R^2^ = 0.17) (Fig. 3c; Table S2). While shoot tissues also differed significantly from each other (R^2^ = 0.15), their microbial community composition were more similar to each other than to those in ovaries (Wilcoxon rank-sum test, p = 0.045), but not significantly more similar than to those of pollen (Wilcoxon rank-sum test, p = 0.128).

### Sources of the seed microbiome

Both sexual and asexual sources made substantial contributions to the seed microbiome, accounting for 30.3% and 23.8%, respectively (Fig. 4a). From the sexual sources, the ovary (22.1%) contributed more than pollen (8.2%; Fig. 4a). From the asexual sources, shoot endophytes at seed maturity contributed a much large fraction of microorganisms to the seed microbiome (21.6%) than the shoot endophytes at bloom (2.2%; Fig. 4a). From the sexual sources, the ovary (22.1%) contributed more than pollen (8.2%; Fig. 4a). Unknown sources accounted for 45.9% of the seed microbiome. Source tracking at the individual tree level showed among-tree variation in source contribution to the seed microbiome (Fig. S1-S3).

**Fig. 4.**
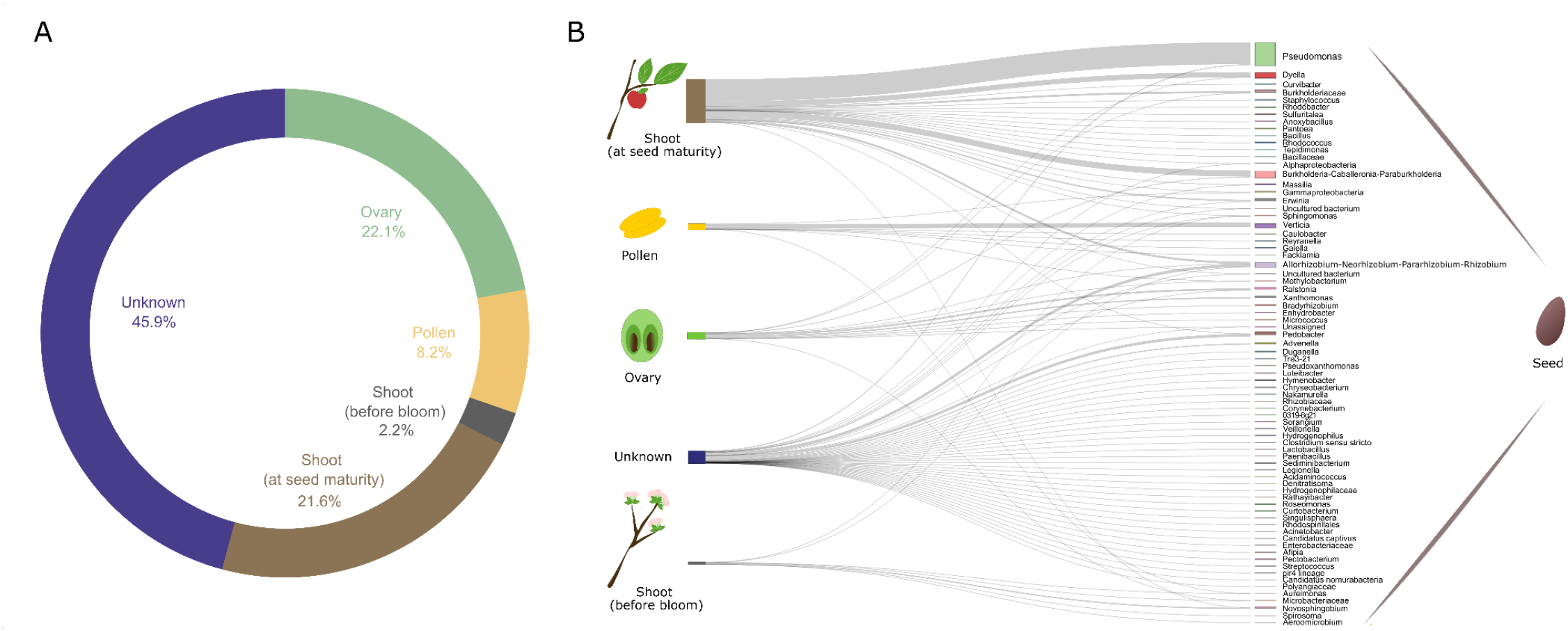
Identification of potential asexual and sexual pathways of microbiome transmission to apple seeds. (A) The donut chart shows the relative contribution of ovary, pollen and shoot endophytes to the seed microbiome, as estimated using fast expectation-maximization for microbial source tracking (SourceTracker). For corresponding standard deviations (SD) and variation between individual trees see Fig. S1. (B) Relative contribution of different sources to the individual genera of the seed microbiome, as estimated using SourceTracker. Each line represents a taxon, with the thickness of the line corresponding to the relative contribution of a given source to the abundance in the seed microbiome. The thickness of each bar on the right shows the overall relative abundance of that genus in the seed microbiome. Taxa are shown at the genus level.

SourceTracker analysis also revealed taxon-specific transmission patterns to seeds (Fig. 4b). *Pseudomonas* and *Burkholderia* colonised the seed microbiome from shoot endophytes collected at seed maturity. In contrast, *Erwinia* and *Dyella* colonised seed from both shoots and unknown sources. Several other taxa, such as *Staphylococcus* and *Rhizobacter*, were primarily associated with pollen or ovaries. Notably, a substantial number of seed-associated taxa, including *Pedobacter* and *Allorhizobium*, colonised seed from unknown sources.

## Discussion

In this study we disentangled the role of sexual (pollen sacs and ovaries) and asexual (shoot endophytes) pathways to seed microbiome. We found that microbial richness, diversity and community composition varied significantly among ovaries, pollen, shoots at bloom, shoots at seed maturity and seed. Sexual and asexual pathways contributed similar proportions to the seed microbiome, with shoots at seed maturity contributing the largest proportion, as followed by the ovary and pollen. We also observed pathway-specific associations for several bacterial taxa: *Pseudomonas* and *Allorhizobium* were primarily linked to shoots, *Rhizobacter* and *Verticia* to pollen, and *Ralstonia* and *Burkholderia* to the ovary. Notably, nearly half of the seed-associated bacterial community could not be traced to the sampled tissues, suggesting that environmental reservoirs such as air or pollinators likely play a substantial role in the origin of the seed microbiome. Overall, these findings indicate the importance of both sexual and asexual pathways in shaping seed microbiomes. Such dual transmission pathways may have evolutionary implications, for example by reinforcing phylosymbiosis.

As expected, we found that shoots at bloom and at seed maturity had higher richness and diversity than the ovary. However, their richness and diversity was not significantly higher than that of pollen. In addition, our hypothesis that shoot endophytes before bloom and at seed maturity are compositionally more like each other than to the ovary and pollen was only partially supported. While microbial communities in shoots were significantly more similar to each other than to ovary, it was not true for pollen. Pollen may serve as microhabitat that support a diverse microbial community due to their high nutrient content, such as sugars, lipids, and proteins (Ambika Manirajan *et al*., 2016; Vannette, 2020). In contrast, the lower diversity observed in the ovary may reflect selective filtering mechanisms, possibly driven by structural barriers or antimicrobial compounds in the ovary (Chesneau *et al*., 2022; Abdelfattah *et al*., 2023). Interestingly, even though ovary had lower bacterial richness than shoots and pollen, they contributed a similar proportion of the seed microbiome. One interesting avenue for future research is to examine when and how microbes disperse between tissues during plant development.

We expected that if sexual pathway played a greater role in microbial transmission, microbes from shoots at bloom might colonize floral organs (ovary and pollen) during their formation, which would later serve as sources for seed microbial communities. This is plausible given that floral tissues originate from vegetative shoots and can share endophytic communities. Alternatively, if asexual transmission dominated, shoots at seed maturity would contribute more to the seed microbiome via vascular connections that facilitate microbial dispersal to the developing seeds. Our findings showed sexual and asexual pathways are contributing nearly equally to microbial transmission to the seed. First, we have demonstrated that sexual tissues together contributed ~30%, with the ovary contributing more (22%) than pollen (8%), though both were consistent sources across tree individuals. The contribution of both male and female reproductive tissues to the seed microbiome may increase microbial diversity and functional redundancy in seeds, particularly under cross-fertilization scenarios (Abdelfattah *et al*., 2023). In parallel, shoot endophytes at seed maturity were the most dominant contributors (~21.6%), indicating that transmission may occur during later stages of seed development. This is biologically plausible and supported by previous studies (Nelson, 2018; War *et al*., 2023), that suggest that as seeds mature alongside the fruit, vascular connections between maternal tissues and the developing seed are likely strongest. Interestingly, while endophytes from shoots collected before bloom had the highest richness and diversity among plant tissues, their contribution to the seed microbiome was minimal compared to the ovaries, pollen and shoots at seed maturity. To better understand the pathways of microbial inheritance, future studies can track the spatial and temporal movement of microbes across tissues, and examine how plant development, immune responses, and tissue connectivity shape transmission pathways.

To date, very few published studies have explored the transmission of microbial communities between plant organs (e.g., ovary, pollen and shoots) and seeds, so the range of bacterial taxa that may colonize seeds via sexual and asexual transmissions is largely unknown. However, a few studies that explored microbial inheritance using watermelons and oaks (Bergmann *et al*., 2025) reported similar results for some of the taxa. For example, showing that, e.g. *Rastonia* and *Bradirhizobium* are likely to be transmitted from the flower to the seed of the plants. As for other taxa, *Pseudomonas*—a dominant seed-associated genus—was primarily traced back to shoot endophytes at seed maturity, with a smaller contribution from the ovary. This aligns with a meta-analysis across 50 plant species, which identified *Pseudomonas* as one of the most prevalent and abundant seed-associated bacteria (Simonin *et al*., 2022). Interestingly, *Pseudomonas* has also been shown to transmit from seeds to the leaves and roots of oak seedlings (Abdelfattah *et al*., 2021), suggesting its central role in both vertical inheritance and subsequent colonization of vegetative tissues. Another abundant taxon, *Burkholderia*, was linked to shoot endophytes at seed maturity. This indicates partial specialization in microbial inheritance, with some taxa primarily using sexual routes, others asexual, and some transmitting by both. A few taxa were shown to be transmitted from pollen. That includes bacteria from *Verticia* genus. Currently, there is no documented knowledge of this bacteria being transmitted from pollen to seeds or being part of the apple seed bacterial community. Thus, its ecological role and implications for seed microbiome establishment represent interesting directions for future research. Among common pathogens of apple trees, *Erwinia* — a causal agent of fire blight disease, was shown: 1) to be present in seeds; and 2) to be mainly transmitted to seeds from unknown sources and to lesser extent from shoots at seed maturity. Several studies have demonstrated the presence of *Erwinia* on stigmas of apple flowers with the possibility that pathogen overwinters on shoots (Piqué *et al*., 2015; Slack *et al*., 2022). Interestingly, diversity of *Enterobacteriaceae* within seeds correlated with higher resistance against pathogenic *Erwinia* species of pumpkin plants in the field (Adam *et al*., 2018), which had implications for microbiome-assisted breeding. Several other abundant taxa, such as a Gram-negative genus *Pedobcater* and *Allorhizobium*, were also shown to be transmitted from unknown sources. *Pedobcater* has been mainly described from the soils (Viana *et al*., 2018), while *Allorhizobium* is primarily known as a nitrogen-fixing symbiont commonly associated with plant root nodules (Kuykendall & Dazzo, 2015). Their presence in apple seeds suggests potential, yet unidentified, transmission pathways or reservoirs within apple trees or surrounding environments, highlighting knowledge gaps in our understanding of microbial inheritance in plant microbiomes.

One interesting result is that nearly half of seed microbiota originated from parent tissues and half from unknown sources (e.g., Abdelfattah et al., 2023). This large proportion of unknown taxa could be explained by unsampled environmental sources of the microbes such as air or mostly likely pollinators. In fact, studies have shown that pollinators can act as vectors of plant-associated microbes, facilitating microbial exchange during flower visits and potentially contributing to seed microbiome assembly (Vannette, 2020; Álvarez-Pérez *et al*., 2024; Cardinale & Schnell, 2024). During foraging, pollinators come into contact with floral organs and nectar, carrying microorganisms on their body surfaces, mouthparts, or digestive tracts. Moreover, the floral microbiome itself is shaped by pollinator identity and behavior, which may influence the microbial communities on reproductive tissues (Prado *et al*., 2020; de Vega *et al*., 2021; Hietaranta *et al*., 2023). As such, insect pollinators may play an important role in transferring microbes from pollen to seeds and offer promising avenue for future research on microbial transmission pathways and pollinator ecosystem services.

Our results also raise the question of whether the assembly follows stochastic or deterministic principles. If you follow the transmission pathways, it looks like a deterministic process, as all organs have very specific microbiomes. The pollen microbiome in particular is unique and ensures the microbial male heritage. In addition, the maternal microbiome changes during fruit ripening, which is also reflected in the seed microbiome, and even shows interesting parallels to the human microbiome (Blaser & Dominguez-Bello, 2016). Overall, our findings highlight the mechanisms behind microbial inheritance in plants, demonstrating that multiple pathways, including sexual (ovary and pollen) and asexual (shoot endophytes), significantly shape the seed microbiome. Understanding these transmission routes has important applied implications for agriculture, as manipulating microbial inheritance could help enhance seedling health, for example disease resistance, through microbiome-assisted breeding.

## Supporting information

see Supporting Information

## Acknowledgements

We are grateful to Jonathan Wilfling and Christian Berg (Botanical Garden) for their support, and to Daniel Höfle for demultiplexing the sequence data.

## Author contribution

AA designed the experiment. AA conducted the sample collection, processing, and AA and DO conducted the molecular work. MF analyzed the data with support from AA. MF wrote the first draft with contributions from AA, AJMT and GB. All authors reviewed the final manuscript.

## Data availability statement

The data that supports the findings of this study, including the ASV table, metadata, R code, are archived in Zenodo repository. These resources can be accessed here via doi: 10.5281/zenodo.15639819 (https://zenodo.org/records/15639819). All amplicon sequencing data generated in this study is deposited on the National Center for Biotechnology Information’s (NCBI) Sequence Read Archive under BioProject accession number PRJNA1308301. They can be accessed through the following link: https://www.ncbi.nlm.nih.gov/bioproject/1308301.

## Supporting Information

**Methods S1** Detailed description of molecular methods and bioinformatics

**Table S1**. Pairwise comparisons of observed bacterial richness and Shannon diversity among different plant tissues: seed, ovary, pollen and shoots before bloom and at seed maturity.

**Table S2**. Pairwise comparisons of bacterial community composition among different plant tissues: seed, ovary, pollen and shoots before bloom and at seed maturity.

**Fig. S1**. Identification of potential asexual and sexual pathways of microbiome transmission to apple seeds using fast expectation-maximization for microbial source tracking (SourceTracker).

**Fig. S2**. Identification of potential asexual and sexual pathways of microbiome transmission to apple seeds using fast expectation-maximization for microbial source tracking (FEAST).

**Fig. S3**. Identification of potential asexual and sexual pathways of microbiome transmission to apple seeds using fast expectation-maximization for microbial source tracking (FEAST).

**Fig. S1.**
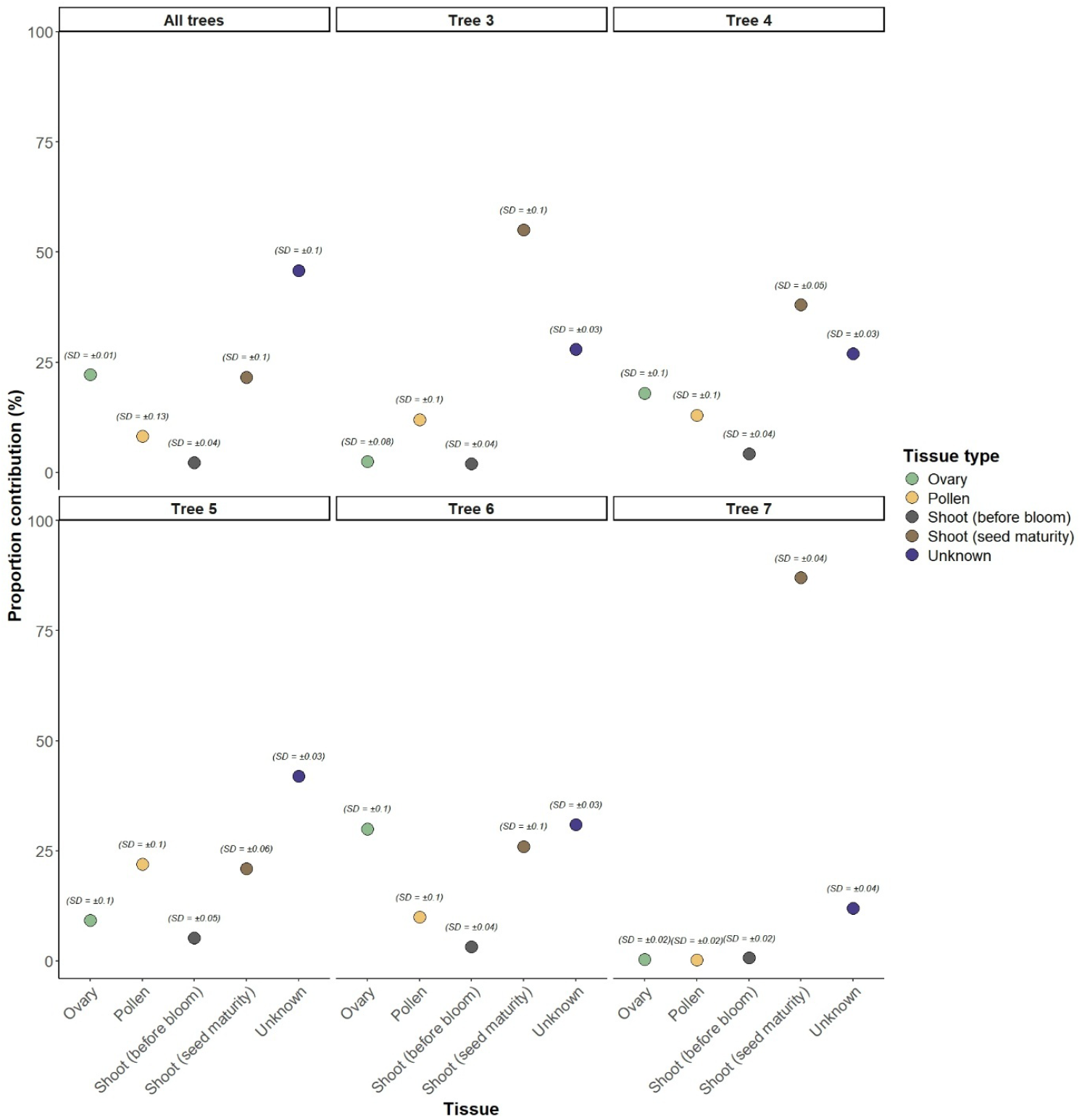
Identification of potential asexual and sexual pathways of microbiome transmission to apple seeds using fast expectation-maximization for microbial source tracking (SourceTracker). Circles represent mean proportion contributions from each tissue type, and the text above denotes the corresponding standard deviation (SD). “Pooled Trees” refers to the aggregated results from all trees, while individual trees show the predicted contribution of each of tissue types to the seed microbiome for each of five trees.

**Fig. S2.**
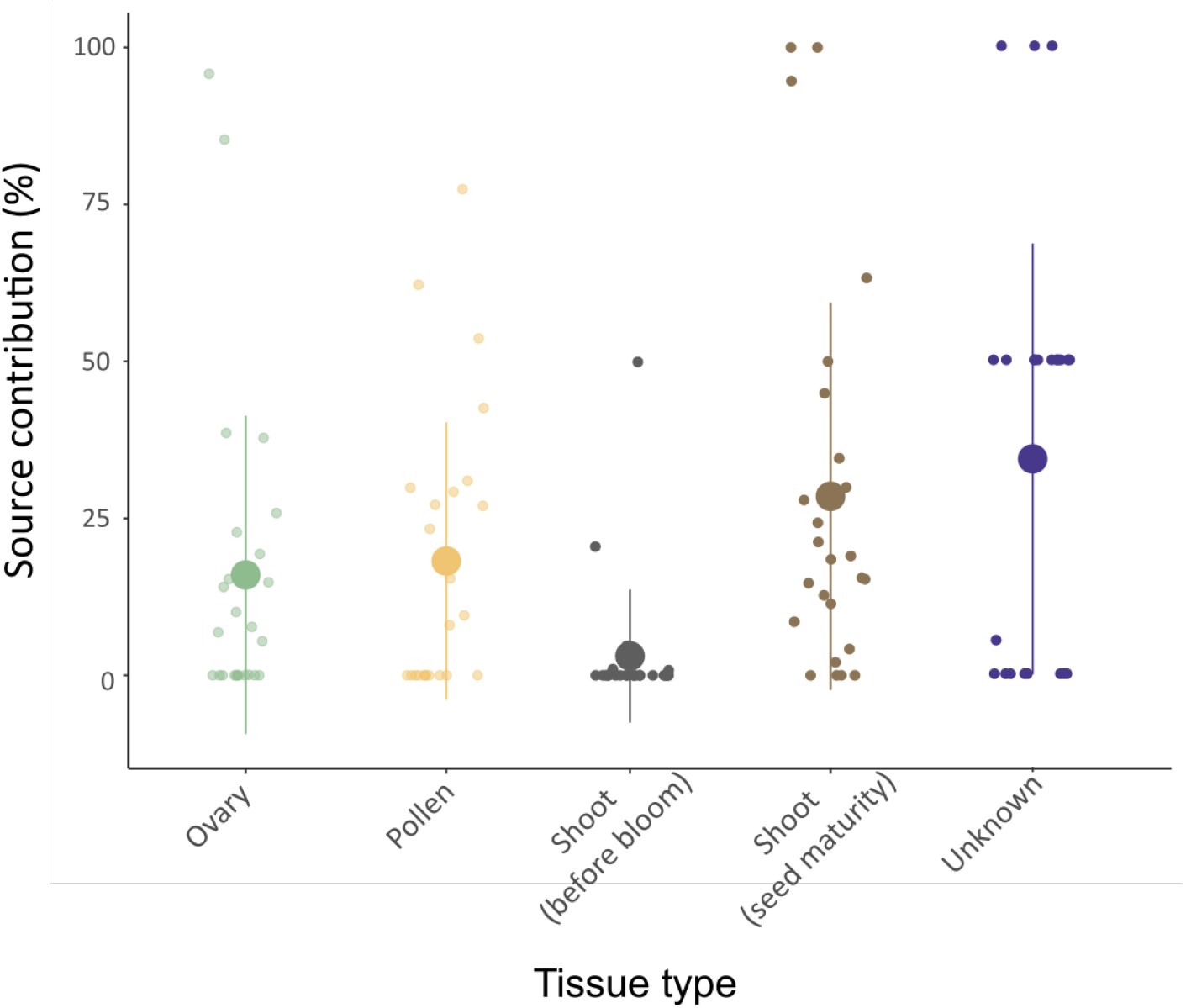
Identification of potential asexual and sexual pathways of microbiome transmission to apple seeds using fast expectation-maximization for microbial source tracking (FEAST). The circle represents the mean values and the error bars represent standard deviations. The small circles represent raw data points, which are horizontally jittered to avoid overlap.

**Fig. S3.**
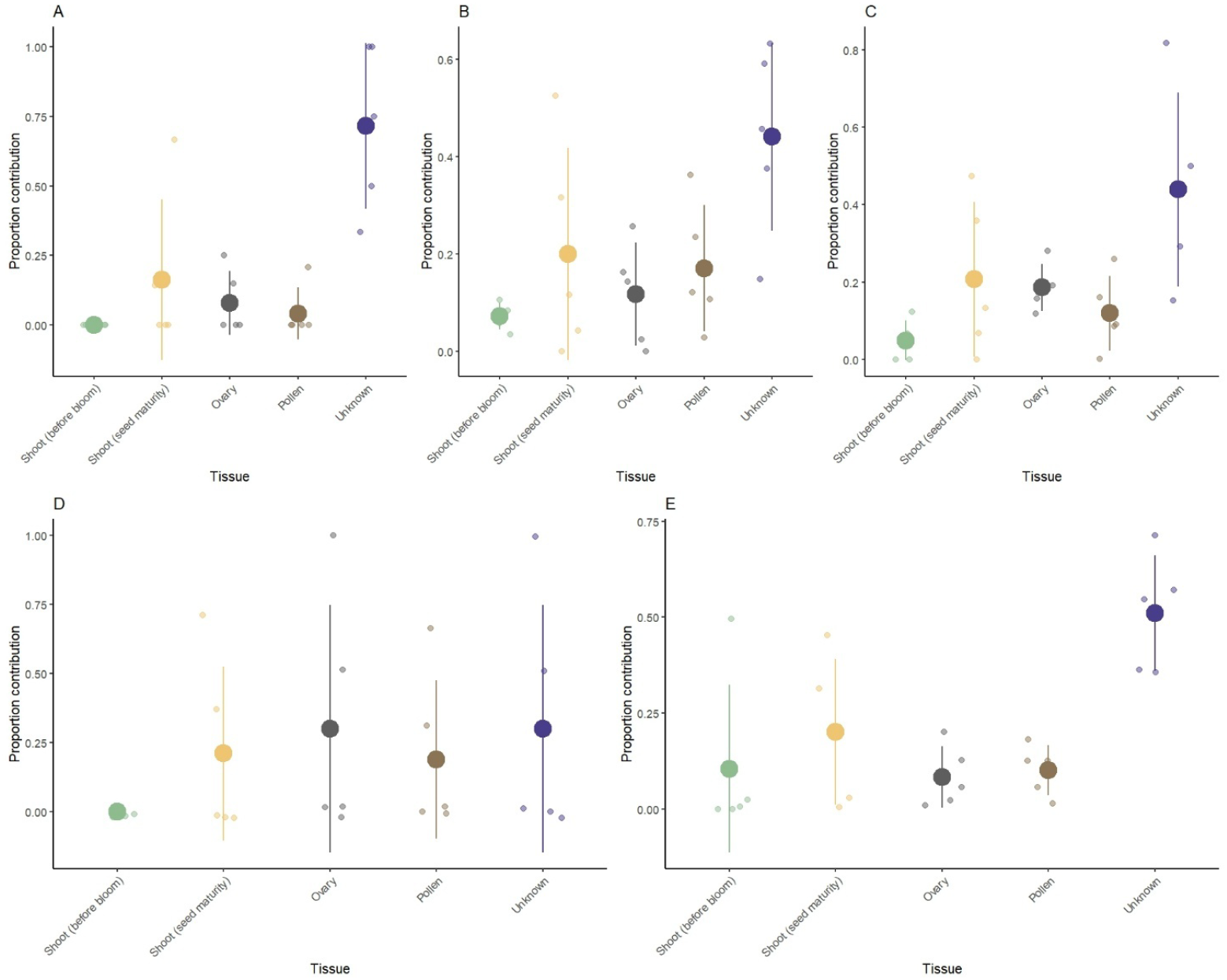
Identification of potential asexual and sexual pathways of microbiome transmission to apple seeds using fast expectation-maximization for microbial source tracking (FEAST). (A-E) Show the predicted contribution of each of tissues types to seed microbiome for each of five sampled trees.

