## Supplementary material for "Plant sexual and asexual contributions to the seed microbiome": see Supporting Information

### Supplementary information

**Methods S1** Detailed description of molecular methods and bioinformatics

DNA was extracted using the FastDNA™ SPIN Kit for Soil (MP Biomedicals). Cell lysis was achieved using a FastPrep Instrument (MP Biomedicals) at 6.0 m/s for 40 seconds. We used 515f and 806r (Walters *et al.*, 2015) to amplify 16S. PCR reactions were run in volume of 30 µL, containing PCR-grade water, 6 µL Taq&Go (MP Biomedicals, Illkirch, France), 0.45 µL of each PNA, 1.2 µL of each primer, and 1 µL template DNA. The PCR protocol consisted of an initial denaturation at 95°C for 5 minutes, followed by 30 cycles of denaturation at 95°C for 30 seconds, a PNA annealing step at 78°C for 5 seconds, primer annealing at 54°C for 30 seconds, and extension at 72°C for 30 seconds, with a final extension at 72°C for 5 minutes.

Raw sequence data were processed using QIIME2 (version 2021.2) (Caporaso *et al.*, 2010; Bolyen *et al.*, 2019). In total, 3216 ASVs and 467,074 sequences passed the filtering controls, with sequence counts ranging from 2 to 127,150 per sample. The pre-rarified taxa table included, on average, 3 ASVs per sample, with counts ranging from 1 to 235. After rarefaction (i.e., 200 reads per sample), the dataset contained 1894 ASVs and 19,400 sequences in total. The number of ASVs detected varied across tissue types: ovaries hosted 261 ASVs, pollen 666 ASVs, shoots at bloom 654 ASVs, and shoots at seed maturity 530 ASVs. The seed microbiome was represented by 330 ASVs.

**Table S1.** Pairwise comparisons of observed bacterial richness and Shannon diversity among different plant tissues: seed, ovary, pollen and shoots before bloom and at seed maturity. Estimates, standard errors (SE), degrees of freedom (df), t-ratios, and adjusted p-values (Tukey method) are shown. Significant comparisons ($p < 0.05$) are in bold.

| **Contrasts** | **Estimate** | **SE** | **df** | **t-ratio** | Adj. **p-value** |
|---|---|---|---|---|---|
| **A) Bacterial richness** | | | | | |
| Ovary vs Pollen | -19.08 | 6.09 | 92 | -3.134 | **0.019** |
| Ovary vs Seed | -6.84 | 5.97 | 92 | -1.145 | 0.782 |
| Ovary vs Shoot (Bloom) | -30.13 | 6.09 | 92 | -4.949 | **<0.001** |
| Ovary vs Shoot (Seed maturity) | -18.52 | 6.03 | 92 | -3.073 | **0.023** |
| Pollen vs Seed | 12.24 | 5.40 | 92 | 2.268 | 0.165 |
| Pollen vs Shoot (Bloom) | -11.05 | 5.52 | 92 | -2.000 | 0.274 |
| Pollen vs Shoot (Seed maturity) | 0.56 | 5.46 | 92 | 0.102 | 1.000 |
| Seed vs Shoot (Bloom) | -23.29 | 5.40 | 92 | -4.315 | **<0.001** |
| Seed vs Shoot (Seed maturity) | -11.69 | 5.33 | 92 | -2.193 | 0.192 |
| Shoot (Bloom) vs Shoot (Seed maturity) | 11.61 | 5.46 | 92 | 2.126 | 0.218 |
| **B) Bacterial diversity** | | | | | |
| Ovary vs Pollen | -0.82 | 0.19 | 92 | -4.335 | **<0.001** |
| Ovary vs Seed | -0.21 | 0.19 | 92 | -1.145 | 0.782 |
| Ovary vs Shoot (Bloom) | -0.95 | 0.19 | 92 | -5.016 | **<0.001** |
| Ovary vs Shoot (Seed maturity) | -0.55 | 0.19 | 92 | -2.916 | **0.035** |
| Pollen vs Seed | 0.61 | 0.17 | 92 | 3.622 | **0.004** |
| Pollen vs Shoot (Bloom) | -0.13 | 0.17 | 92 | -0.750 | 0.944 |
| Pollen vs Shoot (Seed maturity) | 0.27 | 0.17 | 92 | 1.615 | 0.492 |
| Seed vs Shoot (Bloom) | -0.74 | 0.17 | 92 | -4.390 | **<0.001** |
| Seed vs Shoot (Seed maturity) | -0.33 | 0.17 | 92 | -2.014 | 0.268 |
| Shoot (Bloom) vs Shoot (Seed maturity) | 0.40 | 0.17 | 92 | 2.375 | 0.132 |

**Table S2.** Pairwise comparisons of bacterial community composition among different plant tissues: seed, ovary, pollen and shoots before bloom and at seed maturity. Degrees of freedom (df), Sum of Squares (SS), F-value (F), partial coefficient of determination ($R^2$), p-value and adjusted p-value (Adj. p-value) are shown. Significant comparisons ($p < 0.05$) are in bold.

| Contrasts | df | SS | F | $R^2$ | p-value | Adj. p-value |
|---|---|---|---|---|---|---|
| Ovary vs Pollen | 1 | 0.803 | 1.90 | 0.03 | 0.001 | **0.01** |
| Ovary vs Shoot (Bloom) | 1 | 2.823 | 7.34 | 0.12 | 0.001 | **0.01** |
| Ovary vs Seed | 1 | 2.030 | 5.65 | 0.09 | 0.001 | **0.01** |
| Ovary vs Shoot (Seed maturity) | 1 | 1.463 | 3.80 | 0.06 | 0.001 | **0.01** |
| Pollen vs Shoot (Bloom) | 1 | 2.699 | 6.95 | 0.12 | 0.001 | **0.01** |
| Pollen vs Seed | 1 | 1.594 | 4.40 | 0.08 | 0.001 | **0.01** |
| Pollen vs Shoot (Seed maturity) | 1 | 1.202 | 3.09 | 0.06 | 0.001 | **0.01** |
| Shoot (Bloom) vs Seed | 1 | 3.157 | 10.12 | 0.17 | 0.001 | **0.01** |
| Shoot (Bloom) vs Shoot (Seed maturity) | 1 | 2.792 | 8.16 | 0.15 | 0.001 | **0.01** |
| Seed vs Shoot (Seed maturity) | 1 | 1.193 | 3.81 | 0.07 | 0.001 | **0.01** |

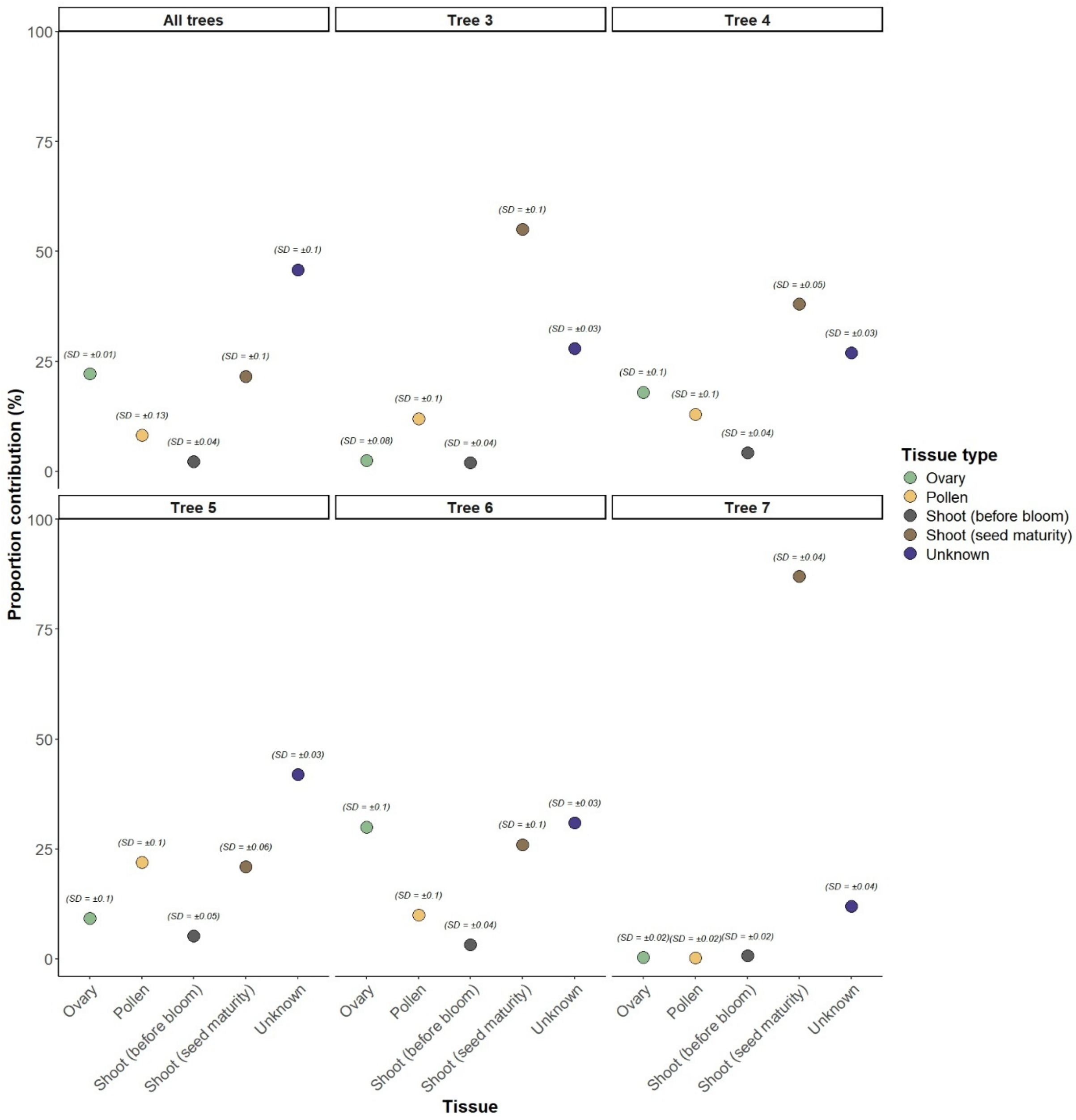


**Fig. S1**. Identification of potential asexual and sexual pathways of microbiome transmission to apple seeds using fast expectation-maximization for microbial source tracking (SourceTracker). Circles represent mean proportion contributions from each tissue type, and the text above denotes the corresponding standard deviation (SD). "Pooled Trees" refers to the aggregated results from all trees, while individual trees show the predicted contribution of each of tissue types to the seed microbiome for each of five trees.

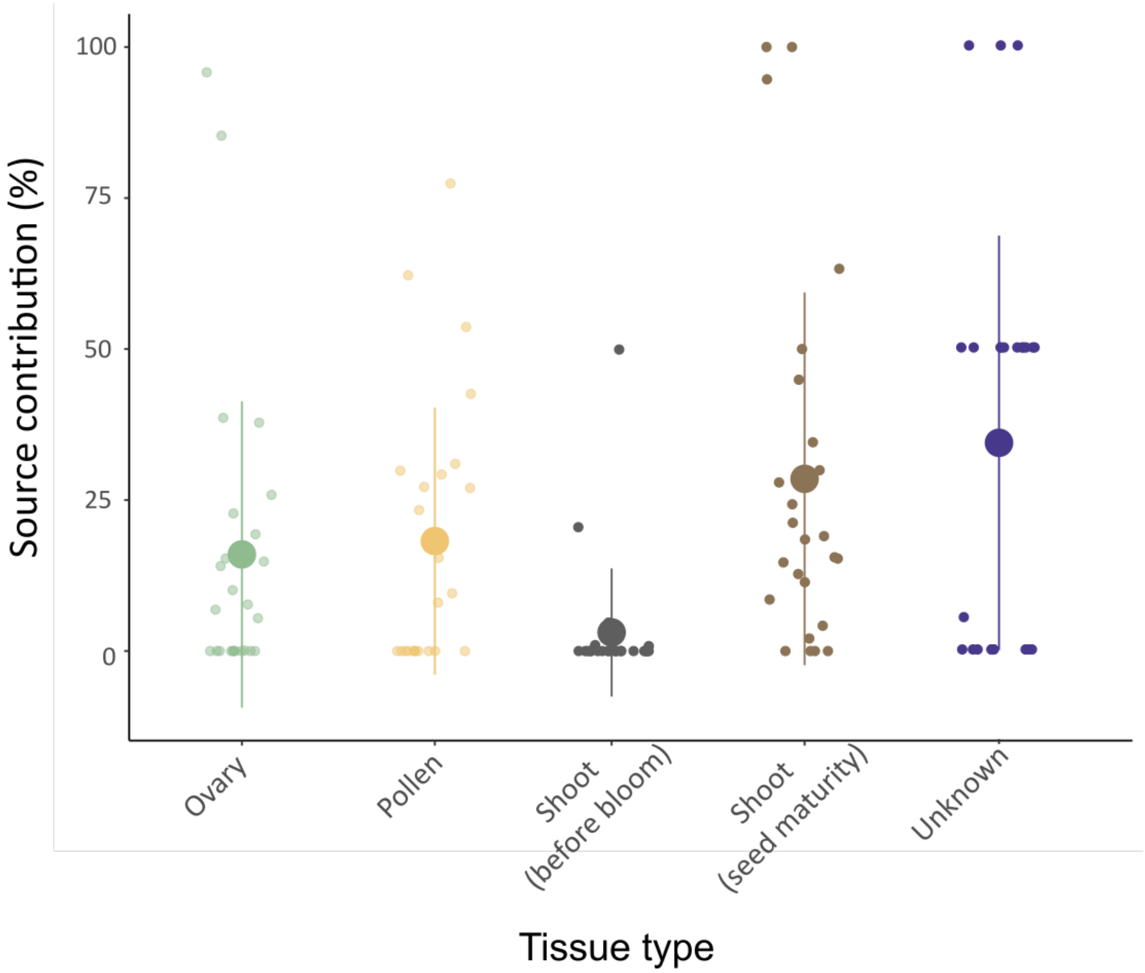


**Fig. S2**. Identification of potential asexual and sexual pathways of microbiome transmission to apple seeds using fast expectation-maximization for microbial source tracking (FEAST). The circle represents the mean values and the error bars represent standard deviations. The small circles represent raw data points, which are horizontally jittered to avoid overlap.

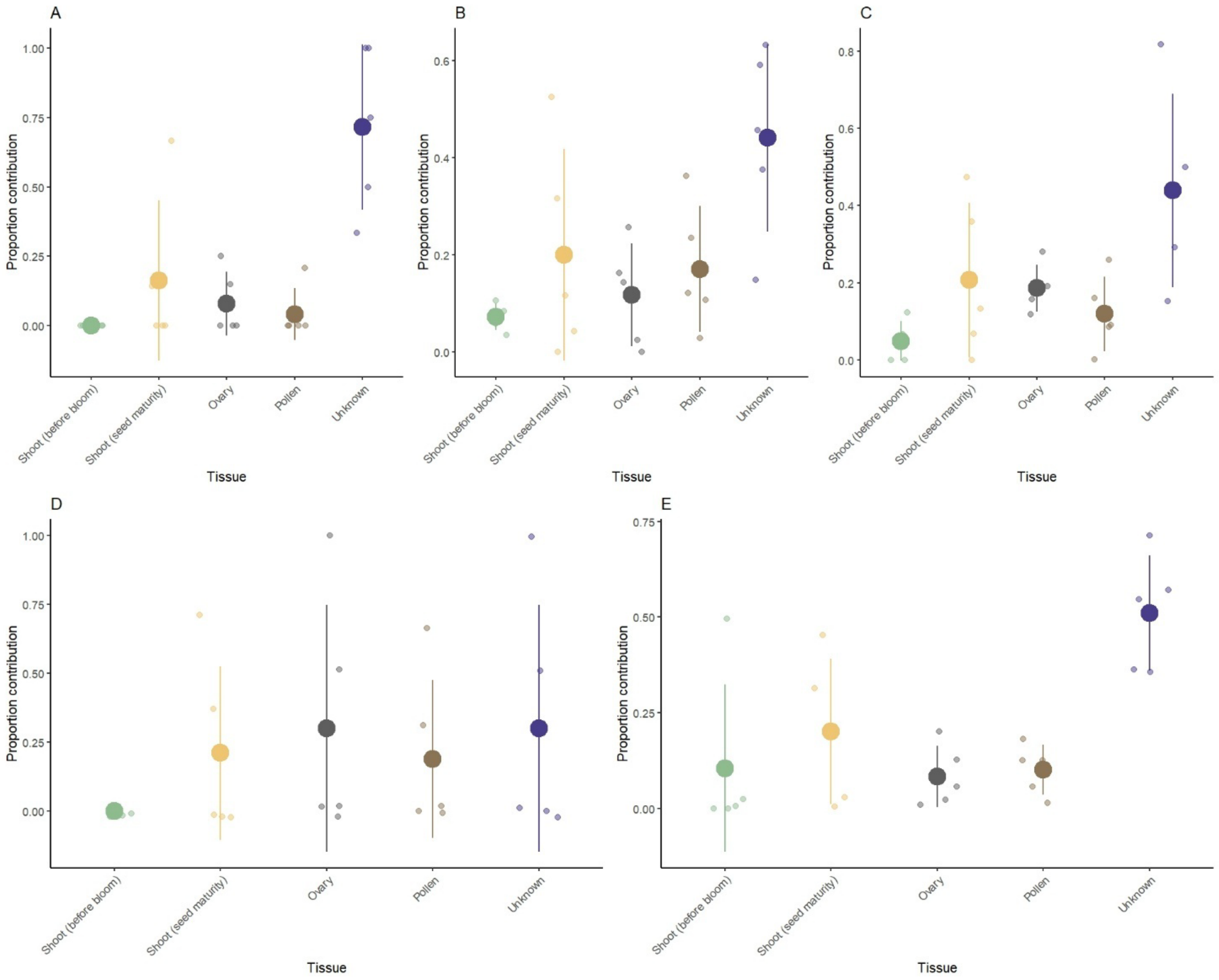


**Fig. S3**. Identification of potential asexual and sexual pathways of microbiome transmission to apple seeds using fast expectation-maximization for microbial source tracking (FEAST). (A-E) Show the predicted contribution of each of tissues types to seed microbiome for each of five sampled trees.
